# ASFV early protein p30 suppresses antiviral type I IFN induction by targeting TRIM21 and RIG-I like receptor signaling adaptor MAVS

**DOI:** 10.64898/2026.03.26.714469

**Authors:** Jiajia Zhang, Hanrong Lv, Jin Ding, Ziyan Sun, Chenglin Chi, Sizhuang Liu, Sen Jiang, Nanhua Chen, Wanglong Zheng, Jianzhong Zhu

## Abstract

African swine fever (ASF) is a highly pathogenic disease caused by the African swine fever virus (ASFV) infection, which can affect pigs of all ages and breeds, posing significant threat to the global pig farming industry. The ASFV p30 protein is an early-expressed viral structural protein; however, its function is not fully understood. In this study, the interaction of viral p30 with host TRIM21 was identified. The ectopic TRIM21 inhibited ASFV replication, while knockdown or knockout of TRIM21 promoted ASFV replication. Further, p30 was found to interact with RIG-I-like receptor (RLR) signaling adaptor MAVS, and during ASFV infection, p30-TRIM21-MAVS interacted with each other. Mechanistically, TRIM21 activated the K27 polyubiquitination of MAVS to induce IRF3 mediated type I interferon (IFN) production, whereas p30 counteracted TRIM21 activated MAVS K27 polyubiquitination to evade RLR signaling mediated antiviral IFN induction. In summary, our study revealed a novel function of ASFV p30, and provided new insights into the immune evasion of ASFV.

## Introduction

African swine fever (ASF) is an acute hemorrhagic viral disease caused by the African swine fever virus (ASFV) infecting domestic pigs and wild boars(1). The mortality rate for pigs infected with the hyperacute and acute types of ASFV is nearly 100%(1). ASFV was first discovered in Kenya in 1921(2), whereas China experienced its first outbreak in 2018(3). Due to the current lack of safe and effective vaccine or specific treatment, it has caused significant economic losses to the global pig industry.

ASFV is the only member of the *Asfarviridae* family and is a large linear double stranded DNA virus(4). The genome of ASFV is vast, encoding over 150 proteins, many of which are functionally unknown or not fully characterized(5). The ASFV p30 protein encoded by viral gene CP204L is an early viral structural protein that can be detected 2-4 h after infection and has good antigenicity(6). Therefore, it is often used as a target protein for vaccine development and serological detection(7). Further, p30 was shown to co-localize with VPS39 and block its binding to HOPS complex, interfering with the function of VPS39 in late endosomal transport(8). p30 can interact with host proteins RPSA, CAPG, DAB2 and ARPC5, which are mainly involved in the endocytic pathway and innate immunity(9). p30 also interacts with RNA-binding proteins CCAR2 and MATR3 to promote ASFV replication(10). Although p30 is an important viral protein, its function has not been fully understood. Therefore, its function in ASFV replication deserves further investigation.

Innate immunity is the body’s first line of defense against pathogen invasion. Upon virus infection, the host encoded PRRs activate the innate immune signaling pathway by recognizing the viral PAMPs(11). As a DNA virus, ASFV is mainly recognized by the innate immune DNA sensing PRR, cGAS, which triggers cGAS-STING-IRF3-IFN antiviral signaling pathway(12). However, ASFV genome DNA is rich in AT regions, so that it can be recognized by DNA directed RNA polymerase III (Pol-III), the transcribed RNA will be subsequently recognized by innate immune RNA sensing RIG-I like receptors (RLRs), activating RIG-I/MDA5-MAVS-IRF3-IFN antiviral signaling pathway(13). Simutanously, ASFV proteins have evolved multiple pathways to evade the cGAS-STING axis mediated antiviral innate immunity(12, 14, 15), as well as RIG-I/MDA5-MAVS axis mediated antiviral innate immunity(13, 16–18).

Mitochondrial antiviral signaling protein (MAVS) is the downstream adaptor of innate immune RLR signaling pathway, and plays a crucial role in host antiviral infection(19). Therefore, MAVS is subjected for tight regulation to ensure optimal protective immune response, and ubiquitination modification is an important regulatory way of post-translational modification of MAVS proteins(20, 21). Triple motif protein 21 (TRIM21) belongs to the E3 ubiquitin ligase family and regulates innate immune pathways via its E3 ligase activity(22). Despite its flexible function, TRIM21 can activate K27 polyubiquitination of MAVS and upregulate IRF3 induced type I IFN, thereby promoting innate immune response to viral infection(23).

In this study, we identified the interaction between p30 and TRIM21, and found the counteraction of ASFV by TRIM21. Further, ASFV p30 interacted with both TRIM21 and MAVS and inhibited TRIM21 mediated K27 ubiquitination of MAVS, suppressing MAVS triggered type I IFN induction. Thus, we have revealed a novel mechanism of ASFV immune evasion, and may provide new insights for the development of antiviral drugs and vaccines.

## Results

### Screening of ASFV p30 interacting proteins and TRIM21 interaction with p30

The ASFV p30 protein persists throughout the entire ASFV replication cycle(10). To identify potential host proteins interacting with ASFV p30 during infection, the porcine primary alveolar macrophages (PAMs) were infected or mock infected with ASFV for 72 h, and the cell samples were collected for ASFV p30 mAb immunoprecipitation. The confirmed immunoprecipitated samples (Fig 1A) were subjected for LC-MS/MS to screen for p30 interacting proteins (Fig 1B). To minimize false positive hits, the p30 interacting proteins in mock cells were used as controls for calibration and those with significantly elevated scores in ASFV infected cells were selected. As such, a total of 189 potentially p30 interacting proteins were identified with ASFV infection (Fig S1). Functional GO and KEGG analysis indicated these proteins are enriched in translation, metabolism, signaling and immune response (Fig S1A and 1B). Given the TRIM21 is involved in innate immune antiviral signaling, it was selected for subsequent investigation (Fig S1C).

**Fig 1.**
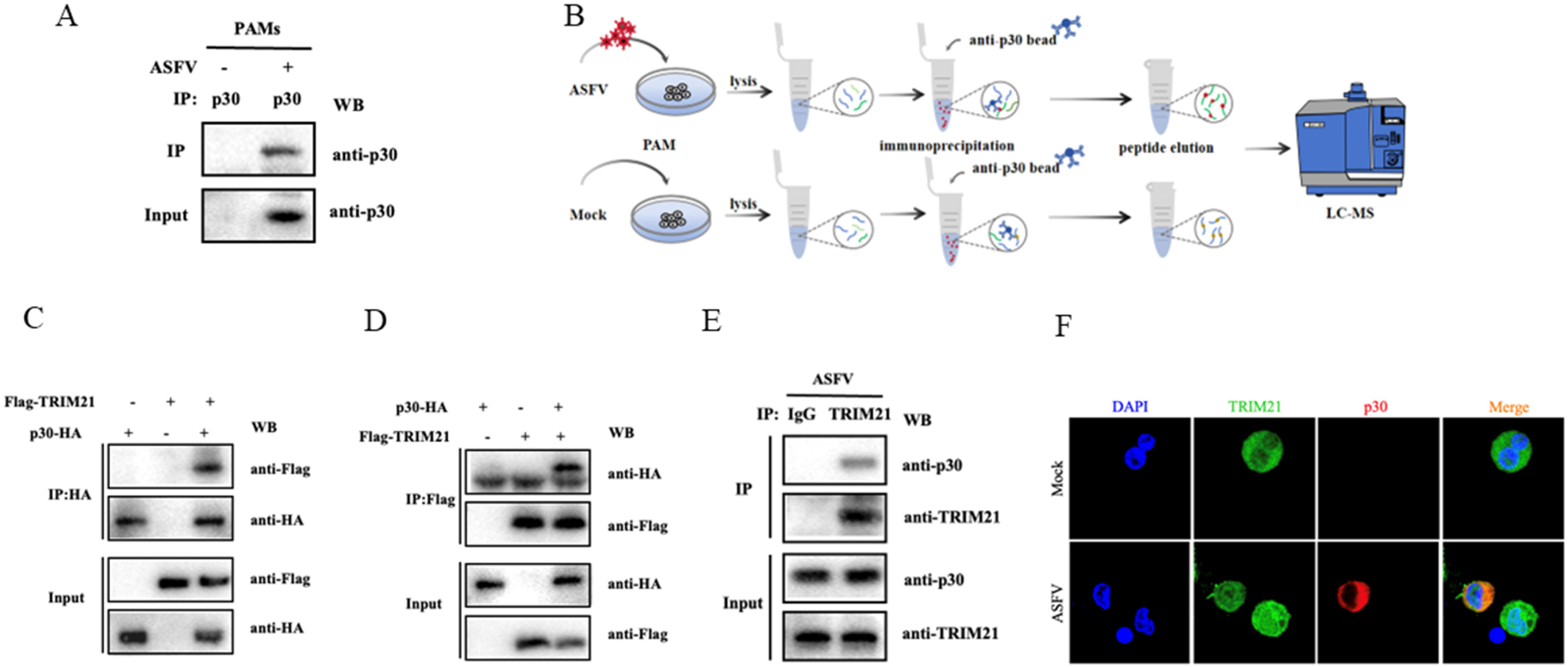
The interaction exists between ASFV p30 and TRIM21. (**A**) The immunoprecipitated p30 protein in ASFV infected PAMs. (**B**) Workflow diagram for ASFV p30 immunoprecipitation and mass spectrometry experiments. (**C** and **D**) 293T cells were co-transfected with p30-HA and Flag-TRIM21 for 24 h. Immunoprecipitation was performed using the specific antibodies, confirming the interaction between exogenous p30 and TRIM21. (**E**) ASFV infected PAMs were harvested 72 h post-infection, and immunoprecipitation was performed using rabbit anti-TRIM21 mAb and control IgG, confirmed the interaction between endogenous p30 and TRIM21. (**F**) PAMs infected with ASFV for 72 h were stained with anti-p30 mouse mAb and anti-TRIM21 rabbit mAb, plus the second antibodies. The stained cells were visualized under confocal microscopy to examine the co-localization of p30 and TRIM21.

Subsequently, the interaction between exogenous p30 and TRIM21 was verified by Co-immunoprecipitation (Co-IP) in 293T cells transfected with p30-HA and Flag-TRIM21. Both Co-IP and reverse Co-IP confirmed the interaction between p30-HA and Flag-TRIM21 (Fig 1C and 1D). The interaction between endogenous p30 and TRIM21 was further examined by Co-IP in ASFV infected PAMs. The results showed that that p30 not only interacts with TRIM21 in ASFV infected PAMs (Fig 1E), but also co-localizes with TRIM21 in the cytoplasm of ASFV infected PAMs (Fig 1F). Together, these results demonstrated that ASFV p30 interacts with host TRIM21 protein.

### TRIM21 is upregulated by ASFV infection and suppresses the replication of ASFV

TRIM21 plays an important role in the natural immune response induced by viruses(24), so we systematically evaluated the role of TRIM21 in host defense against ASFV infection. First, the dynamics of TRIM21 expression during ASFV infection was investigated, porcine PAMs as well as macrophage cell line 3D4/21 cells were infected with ASFV and ASFV-GFP, respectively. Cells were collected at 12 h, 24 h, and 36 h after infections, and TRIM21 expression was detected by Western blotting and RT-qPCR. The Western blotting results showed that the expression of TRIM21 protein increased in both PAMs cells and 3D4/21 cells along the ASFV infections (Fig 2A and 2B). The RT-qPCR results showed that after ASFV infection, the TRIM21 mRNA increased in both PAMs cells and 3D4/21 cells (Fig 2C and 2D). ASFV infection also upregulated the TRIM21 protein and mRNA in both PAMs and 3D4/21, in dose dependent ways (Fig 2E-H). Additionally, TRIM21 protein could be induced by type I IFNα but not type II IFNγ (Fig S2A).

**Fig 2.**
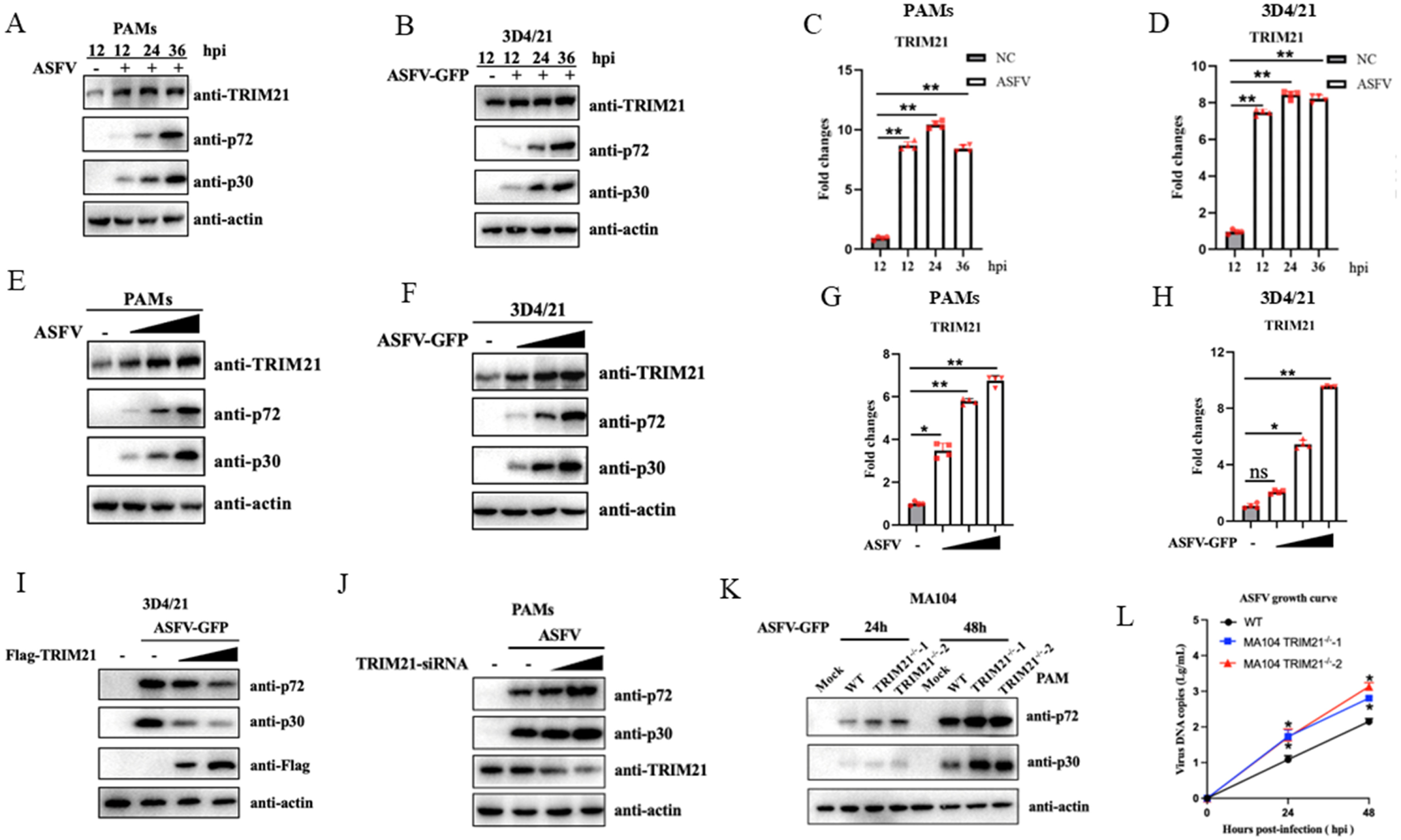
TRIM21 inhibits the ASFV replication in a negative feedback way. (**A**-**D**) PAMs were infected with ASFV (MOI=1), 3D4/21 cells were infected with ASFV-GPF (MOI=1), and cells were collected at 12, 24, or 36 h post-infection. (**E**-**H**) PAMs and 3D4/21 cells were infected with ASFV and ASFV-GFP, accordingly, at different MOIs (0.1, 1 and 2), and cells were collected 48 h post-infection. The expression of TRIM21 was analyzed by Western blotting and RT-qPCR. (**I**) 3D4/21 cells were transfected with Flag-TRIM21 of different doses for 24 h, and infected with ASFV-GFP (MOI = 1). (**J**) PAMs cells were transfected with varying doses of si-TRIM21 or si-NC for 24 h followed by infection with ASFV (MOI=1). Cell samples were collected 48 h post-infection, and the expression of ASFV p72 and p30 proteins was analyzed by Western blotting. (**K** and **L**) TRIM21^-/-^ MA104 cells (clones 1 and 2) and control MA104 cells were infected with ASFV-GFP (MOI=1) for 24 and 48 h, and the viral replication was analyzed by Western blotting and RT-qPCR, respectively.

To further investigate whether TRIM21 is involved in ASFV infection, we conducted ASFV infections in TRIM21 transfected, knockdown and knockout cells (Fig 2I-L). In transfected 3D4/21 cells, Flag-TRIM21 inhibited ASFV p72 and p30 protein expressions in a dose-dependent manner (Fig 2I). In TRIM21 siRNA treated PAMs, the ASFV p72 and p30 protein expressions were upregulated obviously relative to control siRNA treated cells (Fig 2J). Due to the great potential of MA104 cells for ASFV replication(25), two homozygous knockout MA104 cell clones were generated by using CRISPR/Cas9 technology (Fig S2B and 2C). In two TRIM21^-/-^ MA104 cell clones, both ASFV proteins and genome copy were all upregulated significantly relative to infected parent MA104 cells in a time dependent manner (Fig 2K and 2L). Together, these results clearly suggested that TRIM21 plays a negative feedback role in counteracting ASFV replication.

### ASFV p30 targets MAVS to inhibit type I IFN induction pathway

TRIM21 induces the destruction of antibody bound viral particles, followed by DNA or RNA sensors (such as cGAS or RIG-I) recognizing the exposed viral genome in the cytoplasm and activating the immune response(26). To determine whether ASFV p30 affects the cGAS-STING or RIG-I/MDA5-MAVS signaling pathways, 3D4/21 cells were transfected with different doses of p30-HA, and then stimulated with cGAS agonist poly dA:dT, STING agonist 2’3’-cGAMP, and RIG-I/MDA5 agonist poly (I:C), respectively. RT-qPCR results showed that p30-HA could inhibit the mRNA expressions of ISG56 and IFN-β induced by poly (I:C) (Fig 3A), but not by poly dA:dT and 2’3’-cGAMP (Fig 3B and 3C). Similarly, luciferase reporter assay showed that p30 could inhibit the ISRE and IFN-β promoter activities induced by ploy (I:C) in dose-dependent manners (Fig S2D).

**Fig 3.**
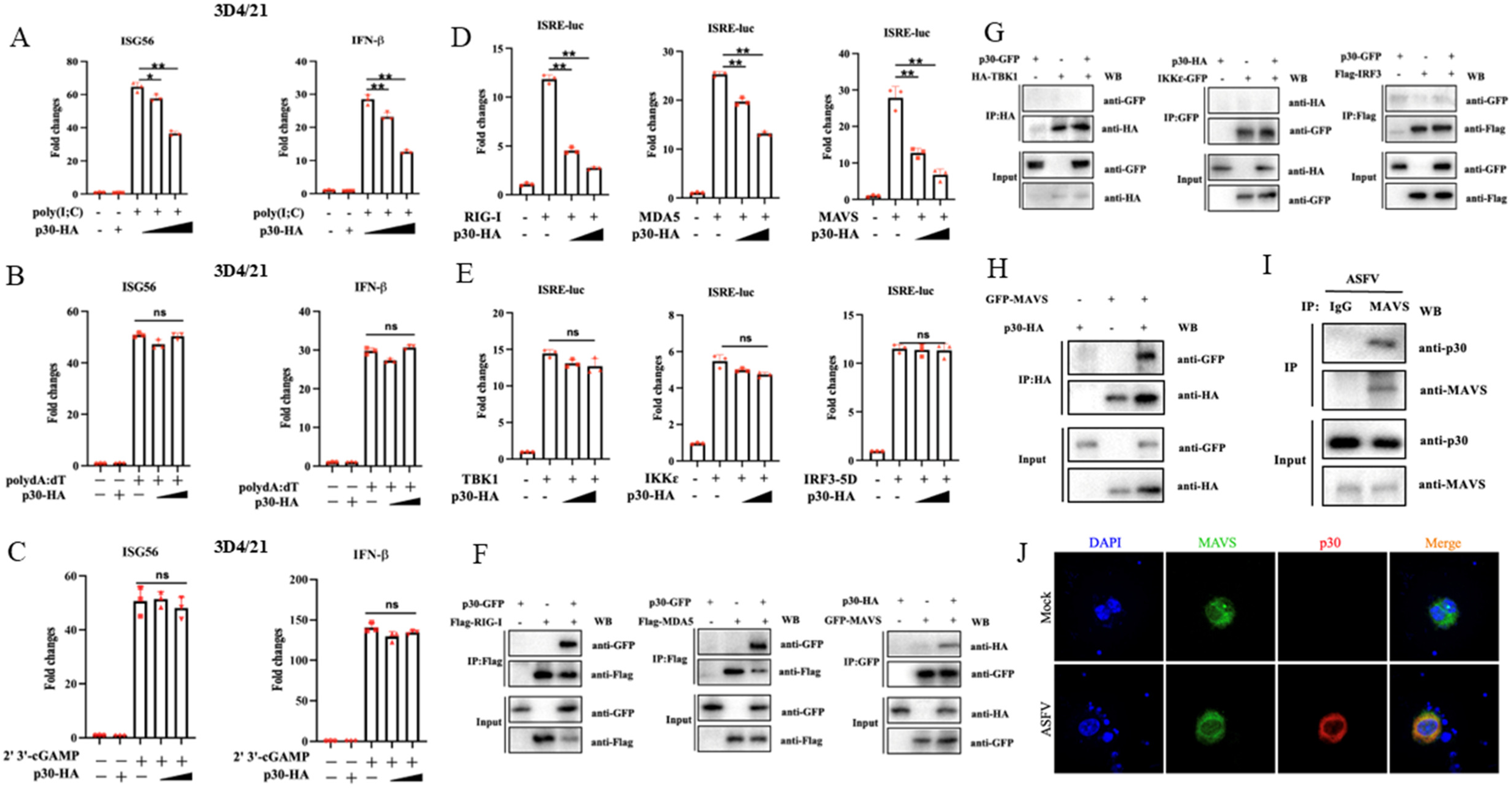
ASFV p30 targets MAVS to inhibit the type I interferon induction signaling. (**A**-**C**) 3D4/21 cells were transfected with different doses of p30-HA for 18 h, followed by transfection stimulation with the 1 μg/mL ploy(I:C) (A), polydA:dT (B) and 2’3’-cGAMP (C) for 12 h. The levels of ISG56 and IFN-β mRNA were detected via RT-qPCR. (**D** and **E**) 293T cells were transfected with RIG-I, MDA5, MAVS, TBK1, IKKε or IRF3-5D, together with different doses of p30-HA and ISRE Fluc and *Renilla* Rluc, to assess the impact of p30 on ISRE promoter activity activated by RIG-I signaling proteins. (**F-H**) 293T cells were transfected with p30 and RIG-I, MDA5, MAVS, TBK1, IKKε or IRF3 for 24 h. Immunoprecipitations (IP) were performed to explore the interactions between p30 and the RIG-I signaling proteins. (**I**) PAMs cells were mock infected or infected with ASFV (MOI = 1) for 72 h, immunoprecipitation was performed using MAVS rabbit mAb to analyze the interaction between endogenous p30 and MAVS. (**J**) PAMs cells were infected with ASFV for 72 h, and confocal microscopy was used to examine the co-localization of p30 and MAVS.

In order to identify the target of ASFV p30 in the RIG-I/MDA5-MAVS signaling pathway, the signaling molecules of RIG-I/MDA5-MAVS signaling pathway, including RIG-I, MDA5, MAVS, TBK1, IKKε, and IRF3-5D(27) were each co-transfected with p30 in 293T cells, and the ISRE promoter activity 24 h post transfection was examined. ASFV p30 was found to inhibit the ISRE promoter activity of RIG-I, MDA5, MAVS, but not of TBK1, IKKε and IRF3-5D (Fig 3D and 3E). In the meantime, the interactions between p30 and RIG-I/MDA5-MAVS signaling proteins were examined in transfected 293T cells by Co-IP assay. The results showed that p30 could interact with RIG-I, MDA5 and MAVS, but not with TBK1, IKKε and IRF3 (Fig 3F and 3G). Based on these results, it was speculated that p30 may target MAVS to inhibit RNA sensing pathway induced type I IFN response. Further, the interaction between p30 and MAVS was confirmed by reverse Co-IP in transfected 293T cells (Fig 3H). Under physiological conditions, the interaction between endogenous p30 and MAVS was confirmed in ASFV infected PAMs by Co-IP assay (Fig 3I), and the co-localization of endogenous p30 and TRIM21 was observed in the cytoplasm of ASFV infected PAMs under confocal microscopy (Fig 3J). Therefore, ASFV p30 was believed to target MAVS to inhibit the antiviral type I IFN induction.

### Protein interactions between ASFV p30, TRIM21 and MAVS

Since ASFV p30 interacted with TRIM21 as well as MAVS, we further investigated the interactions between p30, TRIM21 and MAVS. First, the interaction between MAVS and TRIM21 was examined by Co-IP assay. The results showed that Flag-TRIM21 and GFP-MAVS interacted with each other in transfected in 293T cells (Fig 4A). Second, p30-GFP, Flag-TRIM21 and mCherry-MAVS were all co-transfected into 293T cells, and the Co-IP results showed that there existed mutual interactions between every two proteins (Fig 4B). Third, under physiological conditions, Co-IP showed the MAVS immunoprecipitated both p30 and TRIM21 in ASFV infected MA104 cells, indicating the mutual interactions among these three endogenous proteins (Fig 4C). Finally, to further examine the direct physical interactions among p30, TRIM21 and MAVS, the HA-p30, His-TRIM21 and His-MAVS were purified and the three purified proteins were used in Co-IP experiments (Fig 4D). The results showed that His-TRIM21 and His-MAVS could be pulled down by HA-p30 (Fig 4E and 4F), and His-TRIM21 could be pulled down by His-MAVS (Fig 4G), all convincingly suggesting the direct interactions between p30 and TRIM21, between p30 and MAVS, as well as between TRIM21 and MAVS.

**Fig 4.**
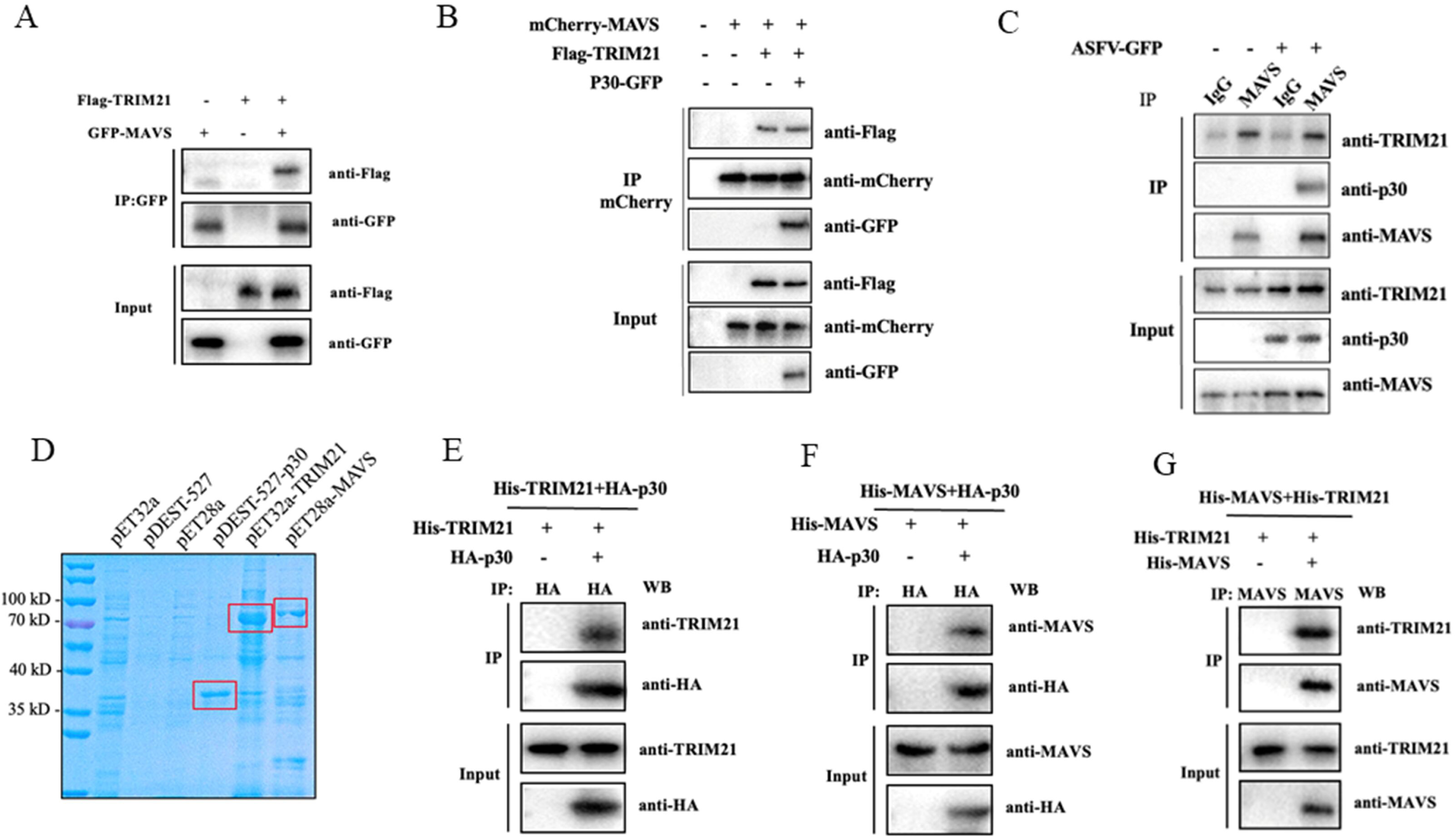
Interactions exist between ASFV p30, TRIM21 and MAVS. (**A**) 293T cells were co-transfected with GFP-MAVS and Flag-TRIM21 for 24 h. Immunoprecipitation was performed using the rabbit anti-GFP mAb to confirm the interaction between exogenous MAVS and TRIM21. (**B**) 293T cells were co-transfected with mCherry-MAVS, Flag-TRIM21, and p30-GFP for 24 h. Immunoprecipitation was performed using rabbit mCherry pAb to confirm the interactions between exogenous MAVS, TRIM21 and p30. (**C**) MA104 cells were infected with ASFV for 72 h. Immunoprecipitation were performed using rabbit anti-MAVS mAb to confirm the interactions between endogenous MAVS, TRIM21 and p30. (**D**) The purified p30, TRIM21 and MAVS recombinant proteins were confirmed by SDS-PAGE and labeled with red boxes. (**E-G**) The interactions between purified proteins p30 and TRIM21, p30 and MAVS, and MAVS and TRIM21 verified through immunoprecipitations.

### Analysis of interaction domains of ASFV p30, TRIM21 and MAVS

We further identified which domains in ASFV p30, TRIM21 and MAVS are critical for their interactions. For ASFV p30, mid-truncation was performed to construct two truncated mutants, representing the N-terminal (1-102 aa) and C-terminal (102-204 aa) regions (Fig 5A). For TRIM21, four truncated mutants were constructed: the RING domain (16-55 aa), the RING domain-deleted mutant (55-474 aa), the PRY/SPRY domain (268-466 aa), and the PRY/SPRY domain-deleted plasmid (1-267 aa) (28) (Fig 5B). For MAVS, three truncated mutants were constructed: the CARD domain (11-78 aa), the PRO domain (104-173 aa), and the C-terminal domain containing TM (173-524 aa) (Fig 5C).

**Fig 5.**
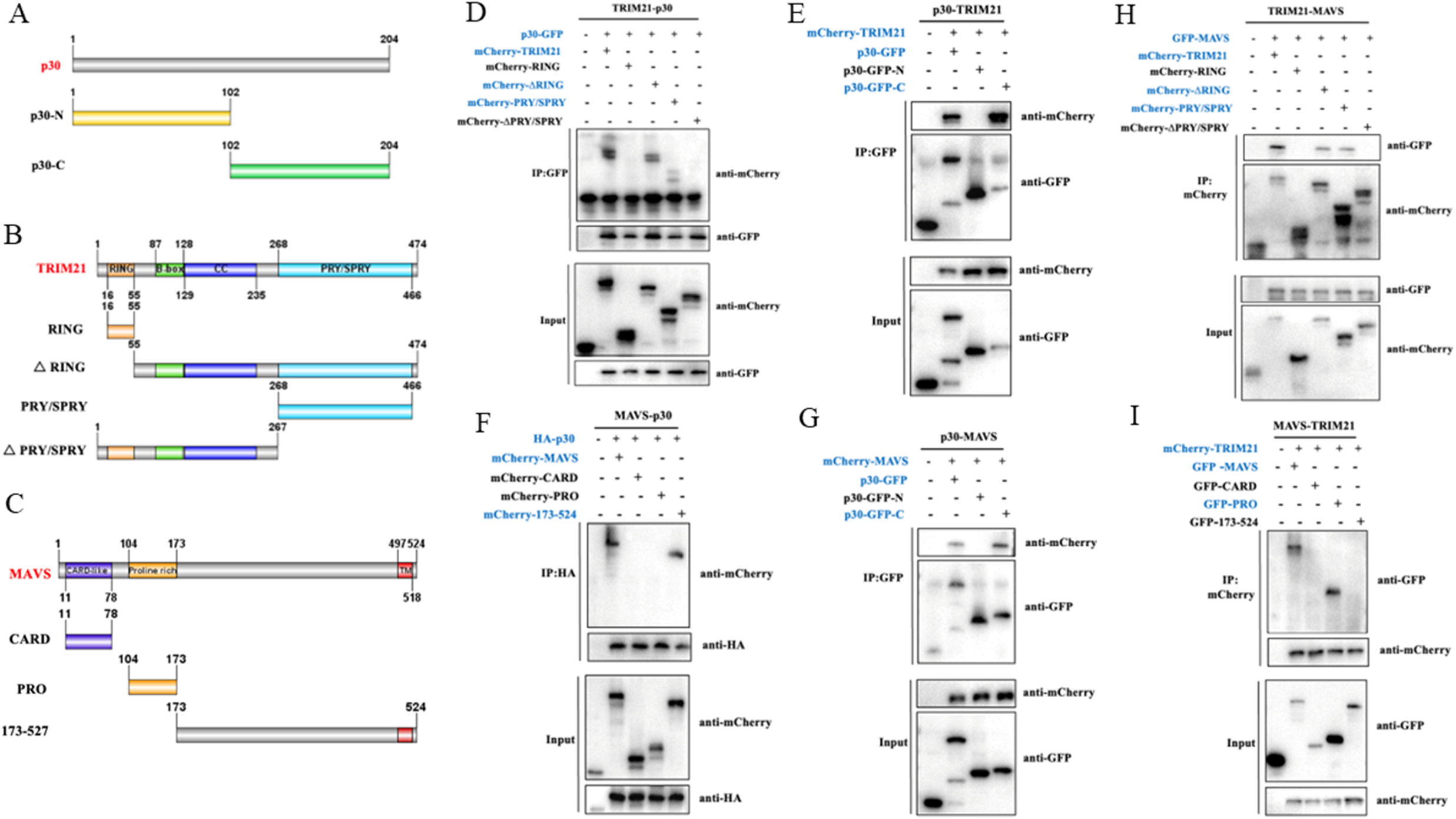
Analysis of responsible domains for ASFV p30, TRIM21, and MAVS interactions. (**A**-**C**) Schematic diagrams of p30, TRIM21 and MAVS truncated proteins. (**D** and **E**) The full-length of p30 interacts with the TRIM21 PRY/SPRY domain (268-466 aa), while the full-length of TRIM21 interacts with the C-terminus of the p30 truncated fragment (102-204 aa) in the immunoprecipitation experiments. (**F** and **G**) The full-length p30 interacts with the TM containing C terminal fragment of MAVS (173-524 aa), while the full-length MAVS interacts with the C-terminus of the p30 truncated fragment (102-204 aa). (**H** and **I**) The full-length of MAVS interacts with the TRIM21 PRY/SPRY domain (268-466 aa), while the full-length of TRIM21 interacts with the MAVS PRO domain (104-173aa). The interacting proteins are labelled in blue color, while those of not interacting proteins are in black color.

For p30 and TRIM21 interaction, the Co-IP results from 293T cells co-transfected with p30 and TRIM21/mutants showed that the p30 can interact with the TRIM21 RING domain-deleted mutant (55-474 aa) and PRY/SPRY domain (268-466 aa) (Fig 5D), suggesting the PRY/SPRY domain is the region of TRIM21 for interacting with p30. Co-IP results from 293T co-transfected with TRIM21 and p30/mutants showed that p30 C-terminal plasmid (102-204 aa) interacts with TRIM21 (Fig 5E). Correspondingly, for p30 and MAVS interaction, the C-terminal domain containing TM (173-524 aa) of MAVS interacts with p30 (Fig 5F), whereas the p30 C-terminal domain (102-204 aa) interacts with MAVS (Fig 5G). In terms of TRIM21 and MAVS interaction, the PRY/SPRY domain is the region of TRIM21 for interacting with MAVS (Fig 5H), whereas the PRO domain (104-173 aa) of MASV interacts with TRIM21 (Fig 5I).

### ASFV p30 inhibits TRIM21 promoted activation of MAVS signaling activity

TRIM21 catalyzes the ubiquitination of K27 in MAVS, further promoting the recruitment of TBK1 to MAVS and activating downstream antiviral signaling pathways(23). We further examined the effect of p30 on MAVS signaling activated type I IFN response in the presence of TRIM21. In transfected 293T cells, the RT-qPCR results showed that ectopic Flag-TRIM21 increased the levels of MAVS activated ISG56 and IFN-β mRNA, whereas p30-GFP could dose dependently inhibit MAVS activated expressions of ISG56 and IFN-β mRNA in the presence of Flag-TRIM21 (Fig 6A). In transfected 3D4/21 cells, RT-qPCR showed that ectopic Flag-TRIM21 promoted poly (I:C) stimulated ISG56 and IFN-β mRNA production, and p30-GFP could dose dependently inhibit poly(I:C) activated expressions of ISG56 and IFN-β mRNA in the presence of Flag-TRIM21 (Fig 6B). The ISRE luciferase assay showed the similar results in transfected 293T cells (Fig 6C).

**Fig 6.**
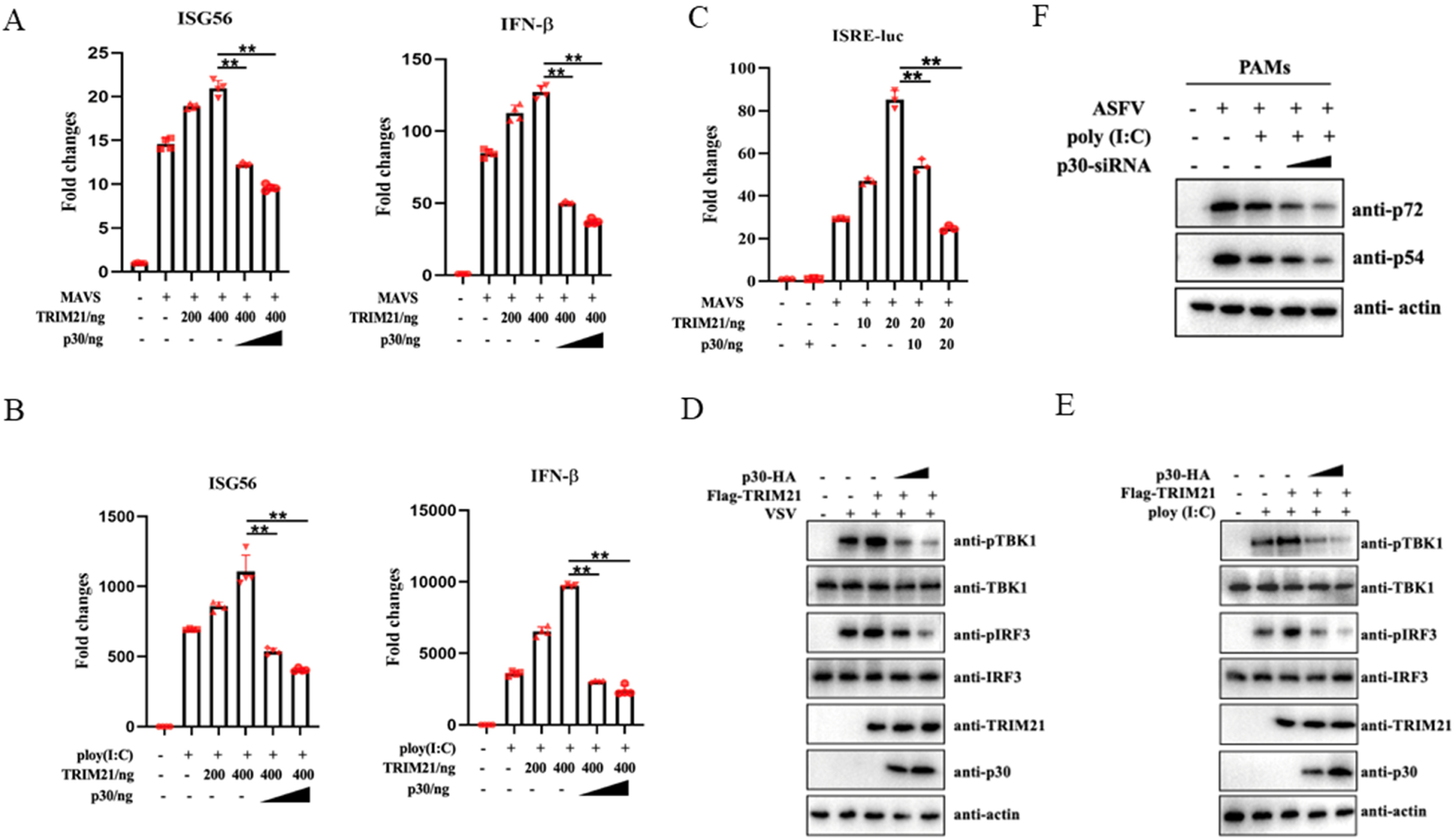
ASFV p30 inhibits TRIM21 upregulated MAVS signaling. (**A**) 293T cells were co-transfected mCherry-MAVS, Flag-TRIM21 and p30-GFP, as indicated, for 24 h. The downstream ISG56 and IFN-β mRNA levels activated by MAVS in the presence of TRIM21 were analyzed using RT-qPCR. (**B**) 3D4/21 cells were co-transfected with Flag-TRIM21 and p30-GFP, as indicated, for 12 h, and stimulated by transfection of poly (I:C) for 12 h. The downstream ISG56 and IFN-β mRNA levels activated by poly (I:C) in the presence of TRIM21 were analyzed using RT-qPCR. (**C**) 293T cells were co-transfected with mCherry-MAVS, Flag-TRIM21 and p30-GFP, plus ISRE Fluc and Rluc reporters, as indicated, for 24 h, and ISRE promoter activity was measured via dual luciferase reporter assay. (**D** and **E**) 3D4/21 cells were co-transfected with Flag-TRIM21 and p30-GFP for 12 h, as indicated. Cells were infected with VSV (D) or stimulated with transfection of poly (I:C) (E), followed by Western blotting for detection of cell signaling activation. (**F**) PAMs were transfected with p30 siRNA or control siRNA for 24 h, stimulated with transfection of poly (I:C) for 6 h, and then infected with ASFV for 48 h. The cell signaling and ASFV replication were detected by Western blotting.

In transfected 3D4/21 cells, Flag-TRIM21 could promote the phosphorylations of TBK1 and IRF3 activated by poly (I:C) (Fig 6D) and by VSV (Fig 6E), and such TBK1 and IRF3 phosphorylations could be inhibited by p30-HA in dose dependent ways (Fig 6D and 6E). To further examine the effect of p30 on poly (I:C) activated anti-ASFV function, PAMs were infected with ASFV, silenced for p30 and stimulated with poly (I:C). The results showed that poly (I: C) stimulation inhibited the expressions of ASFV p72 and p54 proteins, indicating inhibition of ASFV replication, while p30 silence further inhibited ASFV viral replication (Fig 6F). From these results, it could be deduced that p30 can achieve immune escape by inhibiting TRIM21’s promotion of MAVS activation and signaling.

### TRIM21 promotes K27 ubiquitination of MAVS, while ASFV p30 inhibits TRIM21 mediated ubiquitination of MAVS

According to the previous study, TRIM21 can activate MAVS K27 ubiquitination, promoting MAVS activation and antiviral immune response(23). To test such possibility, 293 T cells were first co-transfected with Flag-TRIM21, mCherry-MAVS and Ub-WT plasmids, and the effect of TRIM21 on MAVS ubiquitination was examined. The results showed that TRIM21 could promote MAVS ubiquitination (Fig 7A). When p30-GFP was co-transfected, it could inhibit TRIM21 mediated MAVS ubiquitination (Fig 7B). To further test whether p30 can inhibit TRIM21 mediated K27 ubiquitination of MAVS. 293T cells were transfected with Flag-TRIM21, mCherry-MAVS and Ub-K27 to detect the effect of TRIM21 on MAVS K27 ubiquitination, and the results showed that TRIM21 could promote K27 ubiquitination of MAVS (Fig 7C). When p30-myc was co-transfected, it could inhibit TRIM21 mediated K27 ubiquitination of MAVS (Fig 7D).

**Fig 7.**
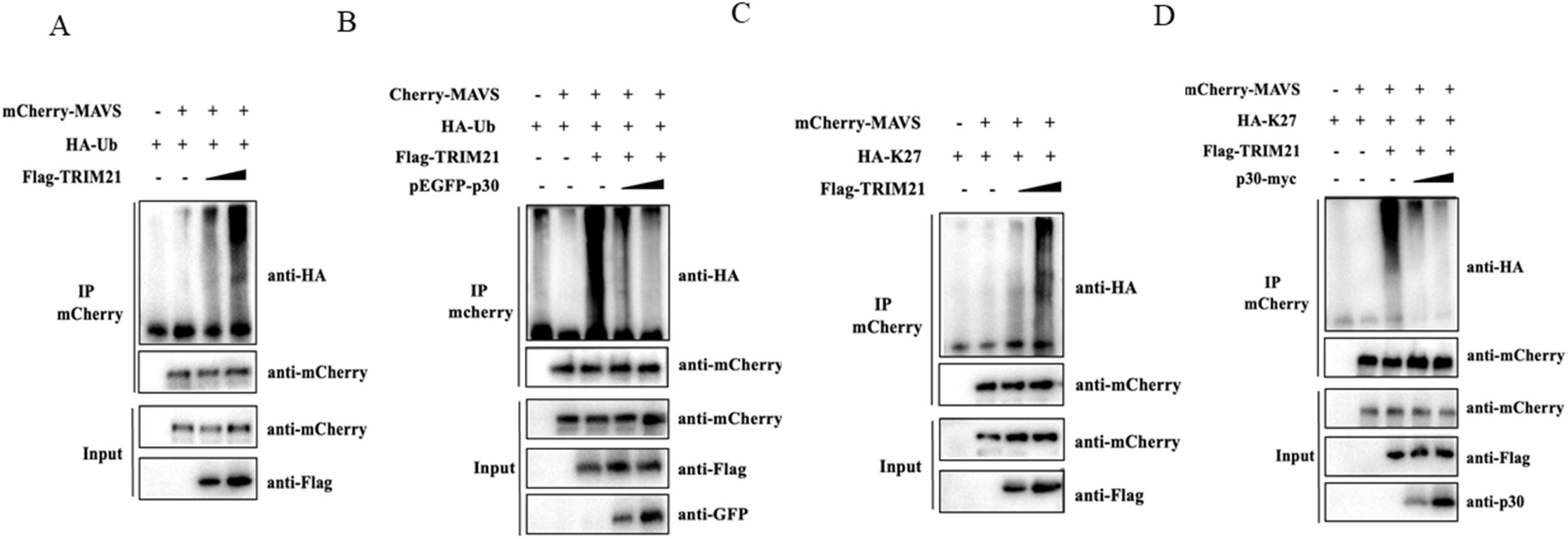
ASFV p30 inhibits TRIM21 mediated ubiquitination of MAVS K27 ubiquitination. (**A**) 293T cells were co-transfected with Flag-TRIM21, mCherry-MAVS and HA-Ub for 24 h and then treated with MG-132 (10 μm) for 6 h. (**B**) 293T cells were co-transfected with Flag-TRIM21, mCherry-MAVS, p30-GFP, and HA-Ub for 24 h and then treated with MG-132 (10 μm) for 6 h. Cell samples were collected and immunoprecipitation were used for analysis of MAVS ubiquitination. (**C**) 293T cells were co-transfected with Flag-TRIM21, mCherry-MAVS and HA-Ub-K27 for 24 h and then treated with MG-132 (10 μm) for 6 h. (**D**) 293T cells were co-transfected with Flag-TRIM21, mCherry-MAVS, p30-myc, and HA-Ub-K27 for 24 h and then treated with MG-132 (10 μm) for 6 h. Cell samples were collected and immunoprecipitation were used for analysis of MAVS K27 ubiquitination.

### ASFV replications in TRIM21^-/-^ MA104 cells complemented with TRIM21, its domains and p30

To explore the key structural domains of TRIM21 in inhibiting ASFV replication. TRIM21^-/-^ MA104 cells were transfected with mCherry-TRIM21, mCherry-TRIM21-RING, and mCherry-TRIM21-PRY/SPRY, respectively, and infected with ASFV-GFP infection. After 48 h infection, fluorescence microscopy, Western blotting and qPCR results demonstrated that the RING domain and PRY/SPRY domain of TRIM21 are essential components, sufficient for TRIM21 to resist ASFV infection (Fig 8A-C). To further determine the promoting effect of TRIM21 on poly (I:C) stimulated anti-ASFV function, and the counteraction by p30, MA104^-/-^ cells were transfected with TRIM21, stimulated with poly (I:C), and infected with ASFV-GFP. After 48 h infection, fluorescence microscopy, Western blotting and qPCR demonstrated that mCherry-TRIM21 further enhances the inhibitory effect of poly (I: C) on ASFV replication, while p30 can reverse this inhibitory effect in a dose-dependent manner (Fig 8D-F). Together, these results clearly suggested that ASFV p30 can achieve immune escape by targeting TRIM21 and MAVS and inhibiting TRIM21 mediated MAVS activation and antiviral signaling.

**Fig 8.**
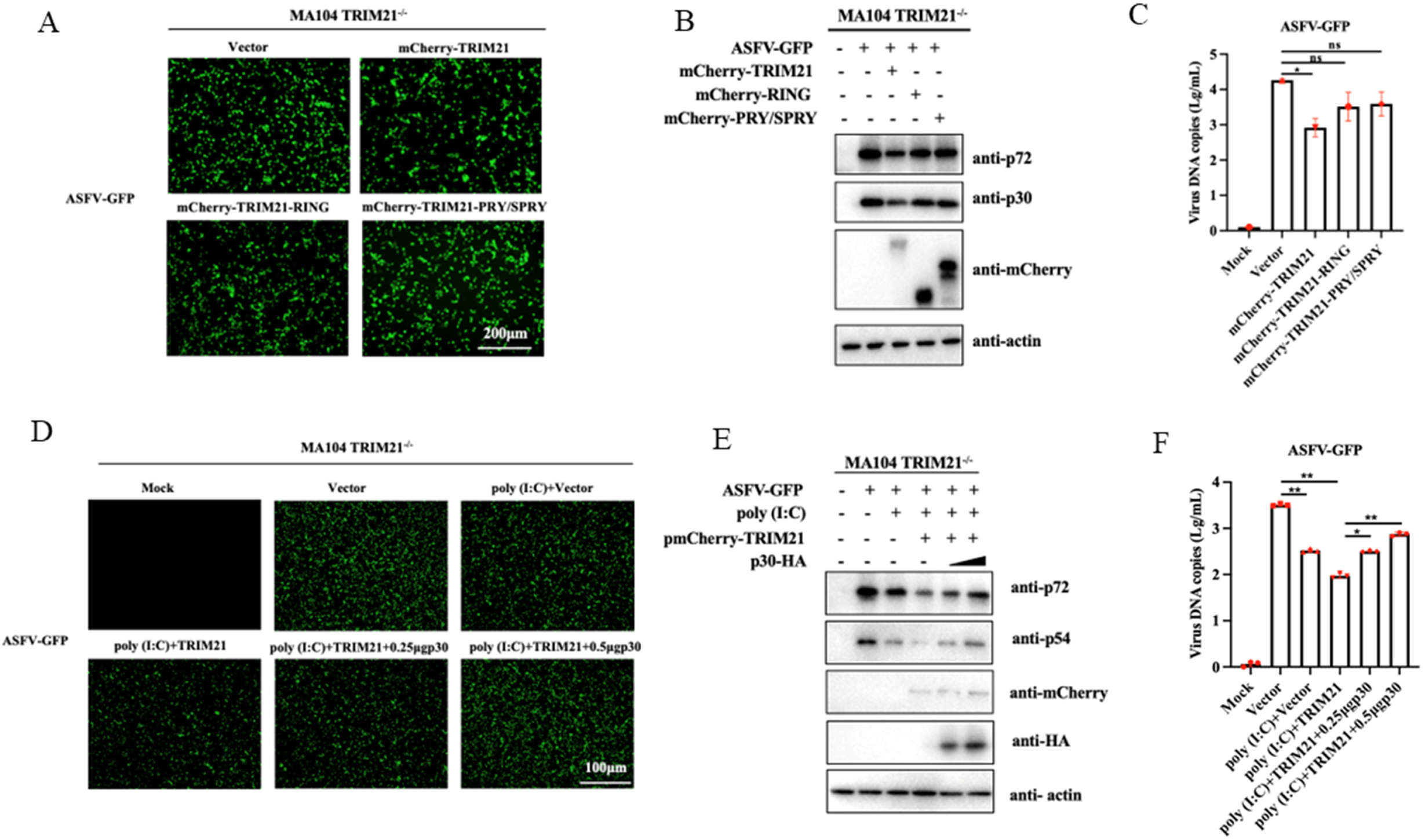
The impacts of TRIM21 and poly (I:C) on ASFV replication. (**A**-**C**) TRIM21^-/-^ MA104 cells were transfected with TRIM21 and its RING, PRY/SPRY domains, respectively, for 24 h, followed with ASFV-GFP infection for 48 h. Virus replications were detected by fluorescence microscopy (A), Western blotting (B) and qPCR (C). (**D**-**F**) TRIM21^-/-^ MA104 cells were co-transfected with TRIM21 and different doses of p30 for 12 h, stimulated with poly (I:C) for 12 h and then infected with ASFV-GFP for 48 h. Virus replication was detected by fluorescence microscopy (A), Western blotting (B) and RT-qPCR (C).

## Discussion

ASFV is a highly pathogenic pathogen that has caused significant economic losses to the global pig farming industry(29); however, due to the large genome and complex immune escape mechanisms, vaccine development for ASFV faces significant challenges. The ASFV p30 protein encoded by the CP204L gene persists throughout the entire virus infection cycle of the virus and is an important immunogenic protein(30). However, understanding of p30 function is still relatively limited at present. In this study, we found that ASFV p30 can inhibit the antiviral type I IFN induction by targeting TRIM21 and suppressing TRIM21 mediated MAVS K27 ubiquitination and MAVS activation, thereby facilitating immune escape and providing favorable conditions for ASFV replication.

As an E3 ubiquitin ligase, TRIM21 can participate in viral infection by directly interacting with viral proteins or regulating immune responses, and may play a bidirectional regulatory role in virus infections(22). Among these situations, viral proteins or host immune molecules can be ubiquitinated by TRIM21 and then degraded by proteasomes, thereby inhibiting viral infection or immune response(22, 31). TRIM21 can also activate K27 polyubiquitination of MAVS and upregulate IRF3 induced type I IFN production, thereby promoting innate immune response to viral infection(23). Regarding ASFV infection, TRIM21 has been utilized to mediate proteasomal degradation of ASFV major capsid protein p72 to resist ASFV infection(32). On the other hand, ASFV can hijack TRIM21 to degrade host immune proteins to achieve immune escape, as exampled by ASFV MGF360-14L and MGF300-2R promoting TRIM21 mediated IRF3 and IKKβ degradation, respectively(33, 34). In our study, we showed that TRIM21 expression can be upregulated by ASFV infection as well as IFNα stimulation, indicating it is an ISG(23). Consequently, TRIM21 can counteract ASFV replication in different cell systems including transfected cells, knockdown cells and knockout cells. Mechanistically, TRIM21 enhances K27 polyubiquitination of MAVS and upregulates IRF3 induced type I IFN, thereby promoting innate immune response against ASFV.

In this study, we found that p30 can affect the RIG-I/MDA5-MAVS signaling pathway, but not the cGAS-STING signaling pathway. The innate RNA sensors RIG-I like receptors (RLRs) mediated pathway have been shown to be involved in ASFV sensing and defense(13), and MAVS acts as the RLR signaling adaptor, subjected for tight regulation(21, 35). In connection with TRIM21, we found ASFV p30 interacts with both TRIM21 and MAVS, and further there exists direct physical interactions between p30, TRIM21 and MAVS. The C-terminal region of p30, PRY/SPRY domain of TRIM21, PRO and C-terminal region of MAVS are the key elements mediating the mutual interactions. In this interaction complex, ASFV p30 is able to suppress TRIM21 mediated K27 ubiquitination of MAVS, thus dampening MAVS activation and its downstream antiviral signaling. The lysine (K) 325 of MAVS is the K27 ubiquitination site catalyzed by TRIM21(23) and K325 site is conserved in porcine MAVS we used in this study, we speculate that p30 may occupy the ubiquitination K325 site and thus block the K27 ubiquitination of MAVS. However, the detailed action of mechanism for p30 inhibition needs to be further investigated.

In summary, our study suggests that TRIM21 is involved in ASFV infection. During viral infection, TRIM21 is upregulated and plays a negative feedback role in counteraction of ASFV infection. ASFV p30 targets both TRIM21 and MAVS, inhibits TRIM21 mediated MAVS K27 ubiquitination and activation, and dampens downstream antiviral signaling (Fig 9). Our study reveals a novel function of ASFV p30 protein and provides new insights into ASFV immune escape.

**Fig 9.**
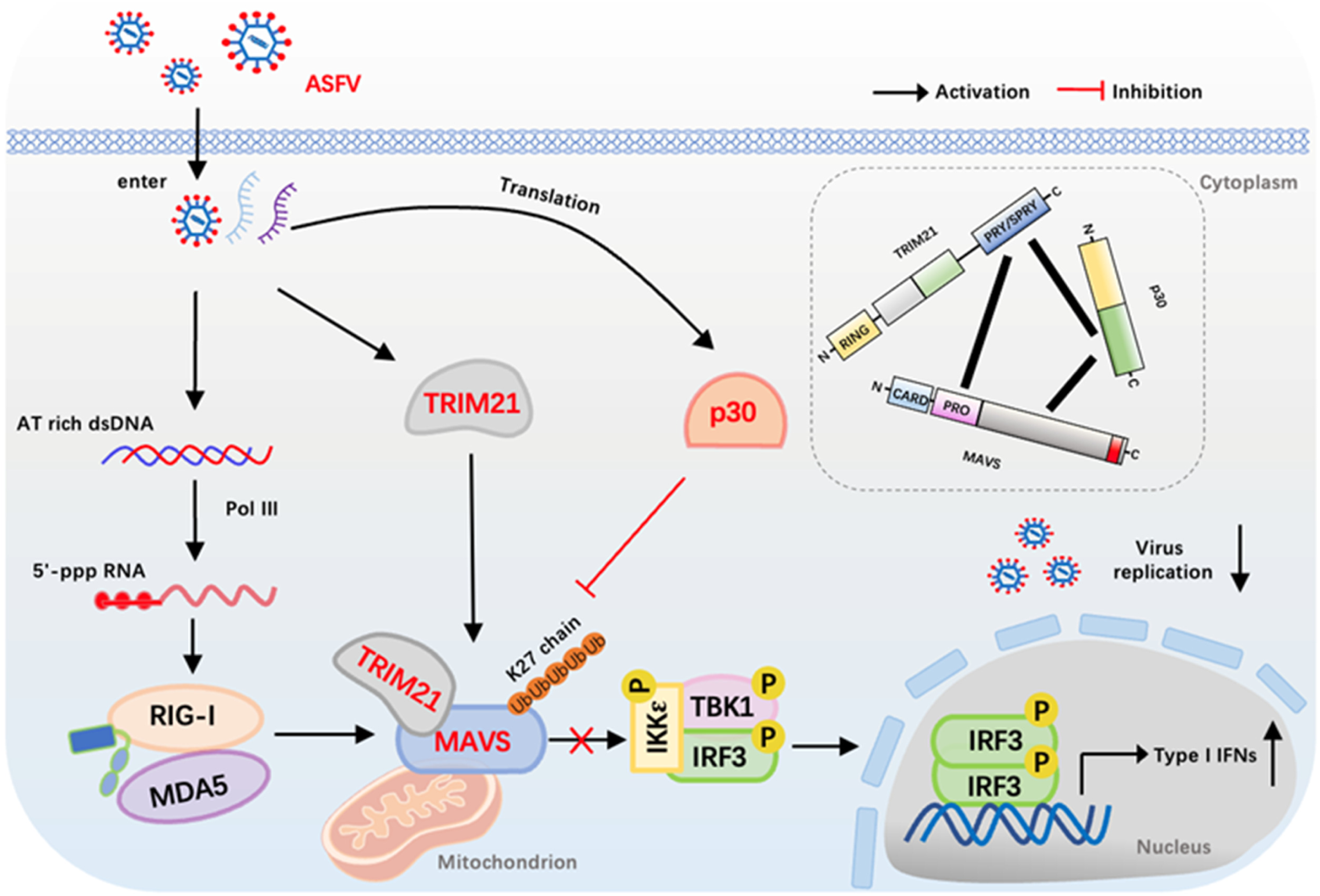
The schematic immune evasion of RIG-I-MAVS antiviral signaling axis by ASFV. Upon ASFV infection, the viral AT rich genomic DNA is recognized by DNA-directed RNA polymerase III (Pol-III), leading to viral RNA sensor RIG-I like receptor (RLR) triggered antiviral innate immune response. Simultaneously, host TRIM21 is induced by infection and acts as the positive regulator of MAVS-IFN activation by RLR sensing. On the other hand, ASFV p30 is expressed, interacts with both TRIM21 and MAVS, blocks the TRIM21 mediated K27 ubiquitination on MAVS, and thus evades the RLR mediated antiviral response.

## Materials and methods

### Cells, cell transfection and viruses

Human embryonic kidney (HEK) 293T (ATCC Cat No: CRL-3216) and MA104 cells (ATCC Cat No: CRL-2378.1) were cultured in DMEM (Hyclone Laboratories, Logan, UT, USA) supplemented with 100 IU/mL of penicillin plus 100 μg/mL streptomycin (Solarbio, Shanghai, China) and 10% fetal bovine serum (FBS) (Vazyme, Nanjing, China). Porcine primary alveolar macrophages (PAMs) were isolated from the fresh lungs of two-month-old piglets of Jiangquhai breed, under the guidance of Experimental Animal Certificate of Yangzhou University under SYXK (JS) 2021-0026. Both porcine PAMs and porcine macrophages cell line 3D4/21 (ATCC Cat No: CRL-2843) were incubated in RPMI 1640 medium (Hyclone Laboratories) containing 100 IU/mL of penicillin plus 100 μg/mL streptomycin and 10% FBS. Cells were grown at 37°C in a 5% CO_2_ humidified incubator. Transfection was performed by using the Lipofectamine 2000 (ThermoFisher Scientific, Shanghai, China) following the manufacturer’s instructions. The ASFV YZ-1 strain (genotype II, GenBank accession ON456300) and MGF100-1R deleted ASFV of YZ-1 (ASFV-GFP) were preserved in the Animal Biosafety Level 3 (ABSL-3) of Yangzhou University approved by the Ministry of Agriculture and Rural Affairs (07140020201109-1).

### Antibodies and reagents

The rabbit anti-TRIM21 mAb, mouse anti-TRIM21 mAb, rabbit anti-MAVS mAb, rabbit anti-IRF3 mAb, rabbit anti-Flag mAb, rabbit anti-HA mAb and rabbit mCherry pAb were all from ProteinTech (Wuhan, China). The rabbit anti-TBK1 mAb (3504S), phosphorylated-TBK1 mAb (p-TBK1, 5483S) were from Cell Signaling Technology (Boston, MA, USA). The rabbit p-IRF3 (Ser385) mAb (MA5-14947) was purchased from ThermoFisher Scientific (Sunnyvale, CA, USA). The mouse anti-FLAG mAb, mouse anti-HA mAb, mouse anti-GFP mAb and, mouse anti-actin mAb were all acquired from Transgen Biotech (Beijing, China). The rabbit anti-GFP mAb was from Beyotime (Shanghai, China). The mouse anti-ASFV p30, p54 and p72 mAbs were all prepared and stored by our laboratory. HRP anti-mouse secondary antibody and HRP anti-rabbit secondary antibody were from BBI (Shanghai, China). Goat Anti-Rabbit IgG H&L Alexa Fluor 594 and Goat Anti-Mouse IgG H&L Alexa Fluor 647 were from Abcam (Shanghai, China). MG-132 (HY-13259) was purchased from MedChemExpress (MCE, Shanghai, China). The Seemless/In-Fusion Cloning (2 × MultiF Seamless Assembly Mix was bought from ABclonal (Wuhan, China). Protein A/G PLUS-Agarose was bought from Santa Cruz Biotechnology (sc-2003, CA, USA). The poly(I:C)-LMW, 2’3’-cGAMP and poly dA:dT were bought from InvivoGen (Hong Kong, China). HiScript® 1st Strand cDNA Synthesis Kit, ChamQ Universal SYBR qPCR Master Mix, 2×Taq Master Mix (Dye plus), 180 kDa prestained protein marker and TransDetect Double-Luciferase Reporter Assay Kit were all from Vazyme Biotech Co., Ltd (Nanjing, China).

### Plasmids and Molecular Cloning

The p30 encoding gene was PCR-amplified from p3×Flag-CMV-7.1-p30 preserved in our laboratory and then cloned into the *EcoR*I and *EcoR*V sites of the vector pCAGGS-HA, the *Bgl*II and *Kpn*I sites of the vector pEGFP-N1, as well as the *EcoR*I and *Bgl*II sites of the vector pCMV-myc, yielding the recombinant plasmids pCAGGS-p30-HA, pEGFP-N1-p30 and pCMV-p30-myc, respectively. The gene encoding porcine TRIM21 (GenBank accession number: NM_001163649.2) was PCR-amplified and cloned into the *Not*I and *Sal*I sites of p3×FLAG-CMV-7.1, generating the recombinant p3×Flag-CMV-7.1-TRIM21. The porcine gene expressing plasmids of pcDNA-DEST47-pRIG-I-Flag, pcDNA-DEST47-pMDA5-Flag, pEGFP-C1-pMAVS, pCAGGS-pTBK1-2HA, pEGFP-C1-pIKKε, p3 × Flag-CMV-7.1-pIRF3, pmCherry-C1-pMAVS, pEGFP-C1-pMAVS-CARD, pEGFP-C1-pMAVS-PRO, pEGFP-C1-pMAVS-173-524, pRK5-HA-Ub, pRK5-HA-Ub-K27 were all previously cloned and preserved in our laboratory. The truncated mutants of p30 were obtained by PCR amplification and cloned into *Bgl*II and *Kpn*I sites of pEGFP-C1 using the 2×MultiF seamless assembly. The porcine TRIM21 and truncated mutants were obtained by PCR amplification and cloned into *Bgl*II and *Kpn*I sites of pmCherry-C1, using the 2×MultiF seamless assembly. For prokaryotic recombinant plasmids, pDEST-527-p30 and pET28a-pMAVS were preserved in our laboratory. Porcine TRIM21 was amplified by PCR and cloned into the *EcoR*I and *Xho*I sites of pET32a vector using 2 × MultiF seamless assembly. The sequences of the cloning PCR primers used are shown in Supplementary Table 1.

### Co-Immunoprecipitation, mass spectrometry and Western Blotting

For co-immunoprecipitation, cells were seeded in a 6-well plate (6-8 × 10^5^ cells/well), collected after transfection or infection, and lysed in 400 μL of lysis buffer for 30 min, followed by centrifugation clearance. An aliquot of 30 μL cell lysate was used as the input control, and the remaining portion was incubated with designated antibodies at 4 °C for 6 h, then 20 μL of G protein agarose beads was added and incubated at 4 °C. After incubation for 3-6 h, the agarose beads were washed four times, then mixed with 2 × SDS sample buffer, and boiled at 100 °C for 10 min. The cleared elution was used for Western blotting. The purified proteins were used as cell lysates for similar co-immunoprecipitation and Western blotting.

For mass spectrometry, PAMs were mock infected or infected with ASFV for 48 h, subjected to immunoprecipitation (IP) with p30 mAb. The IP samples were separated by SDS-PAGE and stained gel slices were excised and analyzed by LC-MS/MS (Novogene Biotechnology Co., Ltd.), enabling simultaneous identification of the target protein and its interacting proteins.

For Western blotting. cell lysates or purified proteins were separated with 6-10% SDS polyacrylamide gel and transferred to PVDF membrane. The membrane was blocked at room temperature with 5% skim milk for 1 h, then incubated with the indicated antibodies overnight in blocking buffer at 4 °C. Next, the membrane was incubated with secondary antibodies for 1 h, and developed for signal with enhanced chemiluminescence (ECL) substrate (Tanon, Shanghai, China).

### RT-qPCR and qPCR

RT-qPCR was performed to detect the expression levels of the target genes. Total RNA was extracted from cells using TRIpure reagent, and reverse transcribed into cDNA by HiScript® 1st Strand cDNA Synthesis Kit. Then target gene expressions were measured by qPCR using SYBR qPCR master mix in StepOne Plus device (Applied Biosystems). The qPCR program was denatured at 95 ℃ for 30 s, followed by 40 cycles at 95 ℃ for 10 s and 60 ℃ for 30 s. TaqMan qPCR was performed to detect ASFV genome copy during replication. The genomic DNA was extracted using DNA mini kit (Cat No: D3121-02, Magen, Guangzhou, China), and then TaqMan qPCR was performed using 2×Taq master mix (Vazyme, Nanjing, China) on StepOne Plus equipment, with the qPCR program of 95 °C for 3 min followed by 40 cycles of 95 °C for 15 s and 60 °C for 1 min. The sequences of the qPCR primers and probe used are shown in Supplementary Table 2.

### Dual luciferase reporting assay

293T or 3D4/21 cells were seeded into 96 well plates (3 × 10^4^ cells/well) and co-transfected with indicated plasmids plus reporter plasmids, ISRE Fluc or IFN-β Fluc, and *Renilla* Rluc with Lipofectamine 2000. Cells were collected at 24 h post transfection and the cell lysates were detected for luciferase activities sequentially by using TransDetect Double Luciferase Reporter Asay Kit. After normalizing Fluc with Rluc, the fold change relative to the control sample was calculated.

### Confocal microscopy

PAMs grown on cover glass in 24 well plates (1-2 × 10^5^ cells/well) were fixed with 4% formaldehyde at RT for 30 min, permeabilized with 0.5% Triton X-100 for 20 min, and then blocked with 5% BSA at room temperature for 30 min. The cells were incubated overnight with primary antibodies (1:500) targeting MAVS, p30, or TRIM21 at 4 ℃, then incubated with secondary antibody (1:500) for 1 h. Finally, the cover glass was counter stained with DAPI, mounted on the glass slide, sealed with nail polish, and visualized under a confocal laser scanning microscope (LSCM, Leica SP8, Solms, Germany).

### RNA silence

The siRNA targeting porcine TRIM21 and ASFV p30 were designed and the siRNAs with the highest scores were selected, synthesized by GeneCreate (Wuhan, China), for subsequent experiments. The siRNA sequences are listed in Supplementary Table 3. According to the manufacturer’s instructions, different doses of siRNA were transfected into PAMs by using Lipofectamine 2000 transfection.

### Preparation of knockout cells by CRISPR-Cas9 editing

In order to obtain MA104 TRIM21 knockout cells, the first exon design of monkey TRIM21 (GenBank: NC_132904) was used as the template for design of CRISPR-Cas9 guide RNAs (gRNAs). Two pairs of high score gRNAs were selected and the annealed gRNAs encoding DNA (Supplementary Table 4) were ligated with the *Bbs*I digested vector pSpCas9(BB)-2A GFP (pX458, Addgene). Subsequently, each gRNA expressing recombinant pX458 was transfected into MA104 cells using Lipofectamine 2000, and GFP positive cells were sorted from the transfected cells using a BD FACSARia III sorting machine. Single MA104 cell clones obtained through limited dilution from sorted GFP expressing cells were screened by PCR using designed primers (Supplementary Table 4). The PCR product was cloned into a T vector using the pClone007 vector kit (TsingKe Biotechnology, Beijing, China). The inserted fragments were subjected to multiplex sequencing, and the insertion and deletion (indel) mutations in the DNA sequence were analyzed. Based on the mutations, two MA104 TRIM21^−/−^ cell clones were obtained (Figure S2B and S2C).

### Purification of ASFV p30, TRIM21, and MAVS proteins

After transforming the prokaryotic expression plasmids pDEST-527-p30, pET32a-TRIM21, and pET28a-MAVS into BL21 competent *E.coli* cells. Positive single colonies were cultured in 15 mL Amp^+^ or Kan^+^ LB liquid medium at 37 ℃ for 3-5 h on a shaker, and IPTG with a final concentration of 1.0 mM was added to the tubes to induce protein expression. The recombinant strain pET32a-TRIM21/BL21 was induced at 37 ℃ for 36 h, the recombinant strain pET28a-MAVS/BL21 was induced at 37 ℃ for 24 h, and the recombinant strain pDEST527-p30/BL21 was induced at 37 ℃ for 12 h. The bacterial precipitates were collected by centrifugation, washed twice with PBS, resuspended with 1mL PBS, and sonicated in on ice for 20 min until the bacterial solution became clear. Subsequently, the MAVS and TRIM21 protein was purified from bacterial lysate supernatant using the His tagged protein purification kit (Beyotime Shanghai, China), whereas ASFV p30 protein was purified from bacterial lysate precipitate and renatured using urea dialysis method(7).

### Statistical analysis

All experiments represent three similar experiments, the bars represent as mean ± SD of different replicates. Experimental data analysis was conducted using GraphPad Prism 10.5.0 software, where *p* < 0.05 was considered statistically significant. In the graphs, “*”, “**”, and “ns” represent *p* < 0.05, *p* < 0.01, and not statistically significant, respectively.

## Figure legends

**Supplementary Fig 1.** Functional analysis of the identified p30 interacting proteins. (**A**) GO analysis plot of the identified interacting proteins. (**B**) KEGG analysis plot of the identified interacting proteins. (**C**) Top 30 potential interacting proteins of ASFV p30, with the TRIM21 marked in red color.

**Supplementary Fig 2.** Induction of TRIM21, TRIM21 knockout cells and effect of p30 on RNA induced signaling. (**A**) PAMs cells were treated with different doses of type I IFNα and type II IFNγ, respectively, for 12 h. Cells were collected and TRIM21 expression was detected by Western blotting. (**B-C**) sequencing results TRIM21^⁻/⁻^ MA104 genomic DNA. The numbers of deleted/inserted bases for each allele are marked on the right side of the sequencing results (“+” indicates insertion and “-” indicates deletion). (**D**) 3D4/21 cells were transfected with p30-GFP for 18 h, and then stimulated by transfection with poly dA:dT and 2’3’-cGAMP for 12 h. The promoter activities of ISRE and IFN-β were detected by dual-luciferase reporter assay.

## Data Availability Statement

The authors confirm that the data supporting the findings of this study are available within the article and its supplementary materials.

## Acknowledgement

This work was partly supported by the National Key Research and Development Program of China (2025YFD1800502-4), the National Natural Science Foundation of China (32473040), the 111 Project under Grant D18007, and A Project Funded by the Priority Academic Program Development of Jiangsu Higher Education Institutions (PAPD). JJ.Z is supported by Research and Practice Innovation Project of Jiangsu Province Graduate Students (SJCX24_2310).

## Author contribution statement

JZ.Z conceived and designed the experiments; JJ.Z, H.L., J.D, Z.S, C.C. and S.L performed the experiments; S.J, N.C. and W.Z provided the resources; JJ.Z and JZ.Z wrote the paper. All authors contributed to the article and approved the submitted version.

